# Dual-matrix 3D culture system as a biomimetic model of epithelial tissues

**DOI:** 10.1101/594549

**Authors:** Diana Bogorodskaya, Joshua S. McLane, Lee A. Ligon

## Abstract

Recent years have seen an unprecedented rise in the use of 3D culture systems, both in fundamental research and in more translational settings such as drug testing and disease modeling. However, 3D cultures often remain underused by cell biology labs, both due to technical difficulties in system setup and inherent drawbacks of many of the common systems. Here we describe an easy to use, inexpensive and rapidly assembled 3D culture system, suitable for generation of both normal polarized epithelial cysts and in-situ tumor spheroids. This system allows for exploration of many questions of normal and cancer cell biology, including morphogenesis, epithelial polarization, cell motility, intra- and intercellular communication, invasion, metastasis, and tumor-stoma interaction. The 3D cultures are made up of a stiffness tunable, dual-matrix model that can incorporate co-culture of multiple cell types. The model allows for increased physiological relevance by mimicking the organization, ligand composition and stiffness present *in-vivo*. The setup allows for a wide spectrum of manipulation, including removing cells from the system for DNA/protein expression, transfection and high-resolution imaging of live or fixed cells.

## INTRODUCTION

### Overview of 3D culture

The first attempts at culturing cells in 3D were made in the early 1980s, in particular with the pioneering work of Mina Bissell and her lab (Bissel 1988). 3D cell culture techniques made incremental progress over the years and slowly gained popularity in the scientific community, with an explosion of interest in the mid-2010s. In recent years, the interest in utilizing 3D cell cultures and organoids as an intermediate platform for drug discovery and toxicity studies has skyrocketed, with multiple techniques developed to bring 3D culture in compatibility with high-throughput systems (Wrzesinski 2015, Nierode 2016). Over the years, multiple advantages of 3D culture systems were highlighted, including increased physiological relevance, ability to dissect the cellular and molecular biology of structures and niches unavailable in 2D cultures, such as epithelial tubes, breast tissue acini and the tumor microenvironment. Currently 3D cell cultures are used to study a broad range of questions, including differentiation, toxicology, tumor biology, morphogenesis and tissue architecture, as well as general cellular properties such as gene or protein expression and cell physiology (Ravi 2015). Multiple culture systems are currently available, ranging from gel-like matrices made from biological extracellular matrix (ECM) components (i.e. Matrigel®, Collagen I), synthetic hydrogel scaffolds (i.e. PEG, PLA) and scaffold-free techniques, such as hanging drop, low-adhesion aggregation or forced flotation (Edmondson, 2015)

The general premise of a 3D culture system is to place an individual cell or a cell aggregate into a 3D matrix, typically a gel or a synthetic scaffold, in which the cells are allowed to grow in all directions. The properties of the matrix are chosen to ensure a physiologically relevant model, with stiffness, ligand presence, and matrix composition typically taken into account. The starting cellular material can also be varied in its level of organization and complexity – from a single cell, which will be allowed to propagate in 3D, to a pre-formed clump of cells, to an organoid with distinct tissue architecture.

However, the widespread adoption of 3D culture systems has been slow due to the technical difficulties of setting up and maintaining the systems as well as the limited toolbox of manipulations and analyses that have been developed to study cells confined in scaffolds. On one side, complicated protocols, high costs of reagents, and long wait times between system setup and ready to use 3D cultures, deter researchers from using 3D setups. On the other side, the limitations of many models, such as high batch-to-batch variability, difficulties manipulating gene expression in 3D, and extracting DNA and protein from the scaffold-confined cells have also contributed to the slow spread of 3D culture use (Katt 2016)

Here we present a new model system with distinct advantages over previous models. First, it is designed to represent the physiological organization of epithelial tissues, with cell aggregates that are surrounded by a model of the basement membrane, which are then further embedded in a collagen-I hydrogel modeling the ECM of connective tissue. In addition, we have developed tools to change the stiffness of this hydrogel to mimic tissues with different mechanical properties. Furthermore, we have developed this model to maximize efficiency and time-to-experiment readiness, and to minimize cost and complexity. And finally, we have refined a set of tools to allow the researcher to manipulate and analyze cells within this 3D culture system, which should expand the utility of this setup. This system can be used to address a wide variety of experimental questions, but here we will use two examples from our work on epithelial morphogenesis and on the tumor microenvironment to illustrate the flexibility of the protocol.

### 3D culture of polarized epithelial cells

Apical-basal polarization is one of the key processes of normal epithelial organization, with defects in polarization often being a hallmark of malignancy (Overeem 2015). When grown in 2D culture, epithelial cells can only achieve partial polarization, although an intermediate 2D/3D culture model, in which the cells are grown on filter inserts, does allow for the growth of polarized cells (Drubin 1996). However, embedding epithelial cells in a gel matrix not only provides conditions for full polarization, but also allows for investigating the cellular and molecular mechanisms implicated in the development of polarized tissues and tissue regeneration (Pollack 1998, Zegers 2003). Madin-Darby Canine Kidney cells (MDCK) are the gold standard model of normal epithelial cells. They have been extensively used in 3D cultures to derive insights regarding mechanisms and signaling behind cell-adhesion and polarization (Capra 2017, Balasubramaniam 2017), as well as morphogenesis and tubule formation (O’Brien 2004, Dolat 2014, Gierke 2012, Boehlke 2013). Other types of cells have been used productively in 3D cultures as well, including primary salivary human stem/progenitor cells (hS/PCs) (Ozdemir 2016), MCF10A breast epithelial cells (Qu 2015), human lung epithelia aggregates (A549), and others (Ravi 2014). However, most of these 3D culture systems involved growing cells in either Matrigel®, collagen-I, or a non-biological hydrogel like alginate or PEG. Matrigel® is a good biochemical model of the basement membrane, but when used as a hydrogel, it loses the structural organization of the sheet-like basement membrane, and collagen-I, while a good biochemical and structural model of the connective tissue ECM, necessitates that the epithelial cells secrete their own basement membrane, which can take a significant amount of time.

### 3D culture of cancer cells

A research area that has significantly benefitted from the development of 3D cell cultures is using cancer cells to model the tumor microenvironment and tumor-stoma interactions. Collagen-I hydrogels were one of the earliest 3D methods used to study the response of cancer cells to the extracellular matrix (ECM) (Richards et al., 1983). Many reports have been published examining the behavior of tumor cells in collagen-I or other, more specialized or defined, 3D matrices, but few if any of these models were representative of the organization of pre-metastatic tumors. Cells from a tumor *in situ* are exposed to a different ECM than metastatic cells; specifically, a tumor *in situ* is encapsulated within a basement membrane (BM), while a metastatic cell has left the tumor site and invaded into the connective tissue stroma. The basement membrane and the stroma are composed of different constituent components. BM is primarily made up of type IV collagen, laminin, and heparan-sulphate proteoglycans (Kalluri, 2003), while the stroma is largely made up of type I collagen and elastin (Culav et al., 1999). The physical characteristics of the two compartments differ as well; a solid tumor is typically much stiffer than the surrounding stroma (Paszek et al., 2005; Levental et al., 2009). Therefore, a biomimetic model of a tumor *in situ* should consider the composition and physical properties of both compartments and the organization of the two compartments relative to each other.

## MATERIALS

### Reagents

- Acetic acid (glacial; C_2_H_4_O_2_)
- Bovine serum albumin (BPA; product BP9703, Fisher Scientific)
- Collagenase (product 02195109, MP Biomedicals)
- Collagen-I (product 150026, MP Biomedicals)
- Dimethyl sulfoxide (DMSO; product D2650, Sigma-Aldrich)
- Distilled water (ddH2O)
- DMEM (product 10-013-CV, Corning)
- Ethylene glycol-bis(succinic acid N-hydroxysuccinimide ester) aka PEG-diNHS (product E3257, Sigma-Aldrich)
- Glassware (bottles with caps: 100 mL, 1 L)
- Goat serum (product G6767, Sigma-Aldrich)
- Growth factor reduced Matrigel® (product 354230, Corning)
- Magnetic stirrer and stir bar
- OPTI-MEM I (product 31985-070, Life Technologies)
- Paraformaldehyde solution (product 18814, Polysciences Inc)
- SlowFade® Diamond (ThermoFisher Scientific)
- Sodium azide (NaN_3_; product 190385000, Acros Organics)
- Sodium chloride (NaCl; product S5886, Sigma-Aldrich)
- Sodium phosphate dibasic (Na_2_HPO_4_; product S5136, Sigma-Aldrich)
- Sodium phosphate monobasic (NaH_2_PO_4_; product 71505, Sigma-Aldrich)
- Sodium pyruvate (prodct 11360070
- Triton X-100 (product BP151, Fisher Scientific)
- µ-Slide 8-well Glass Bottom chamber slide (product 80827, Ibidi)
- 1.5 mL disposable microcentrifuge tubes
- 4D-Nucleofector® X Kit (product V4XC-1032, Lonza)
- 10 cm cell culture dishes (product 172958, Thermo Scientific)
- 15 mL disposable conical tube with cap (product 352097, Becton Dickinson)
- 35 mm tissue culture dishes (product10861-586, VWR)
- 35 mm glass bottom microwell dish (product PG5G-1.5014-C)
- 50 mL disposable conical tube with cap (product 82018-050, VWR)
- 60 mm tissue culture dishes (product 10062-890, VWR)
- 96-well round bottom ultra-low attachment microplates (product 7007, Corning)
- 100 mm tissue culture dishes (product 10861-594, VWR)

### Equipment

- Class II microbiological safety cabinet
- CO_2_ cell culture incubator
- Fluorescent and light microscope (model DMI4000 B, Leica Microsystems)
- Forceps
- Hemocytometer
- Nutating shaker (model 117, TCS Scientific)
- Kimwipes (product S-8115, Kimberly-Clark)
- Pipettes with non-sterile and sterile plastic tips (P2, P20, P200 and P1000)
- Rotational shaker (product 6780-FP, Corning)
- Sterile spatula
- 4D-Nucleofector (units AAF-1002B, AAF-1002X, Lonza)

## METHOD AND PROTOCOL

### Cell culture

MDCK cells (product ATCC CCL34, American Type Culture Collection, Manassas, VA) were cultured in DMEM high glucose (Corning) supplemented with 10% fetal bovine serum (VWR, Radnor, PA), 1% penicillin/streptomycin (Corning, Corning NY) at 37 °C, 5 % CO_2_. MDA-MB-231 human mammary epithelial tumor cells (ATCC HTB-26, American Type Culture Collection, Manassas, VA) were cultured in DMEM (Mediatech, Manassas, VA) supplemented with 5% fetal bovine serum (Atlanta Biologicals, Lawrenceville, GA), L-glutamine (Mediatech, Manassas, VA) and penicillin/streptomycin (Mediatech, Manassas, VA) under 5% CO_2_. For both cell lines, cells from passage 5 – 25 were used.

### Preparation of polarized epithelial spheroids

An overview of the epithelial spheroid model production protocol is depicted in Figure 1. Briefly, MDCK cells are harvested through trypsinization and counted. Pipette 250,000 cells in 1 mL of fresh culture media into a 15 mL conical tube. Then add 500 µL of 3 mg / mL growth factor-reduced Matrigel® diluted in OPTI-MEM I media to the cell solution. This results in a final concentration of 1 mg/mL Matrigel®, which is below the concentration necessary for gelation. This allows the basement membrane extracellular matrix (ECM) components in Matrigel® to adsorb to the cell surface and jumpstart the formation of the basement membrane. The tube should then be placed on its side in the incubator with the cap partially open to allow air exchange, and incubated overnight. This will allow the cells to coalesce into small cell clumps. Most epithelial cells will preferentially adhere to one another rather than the non-tissue culture plastic of the tube. If the cells adhere to the tube, try tubes from different manufacturers.

**Figure 1:**
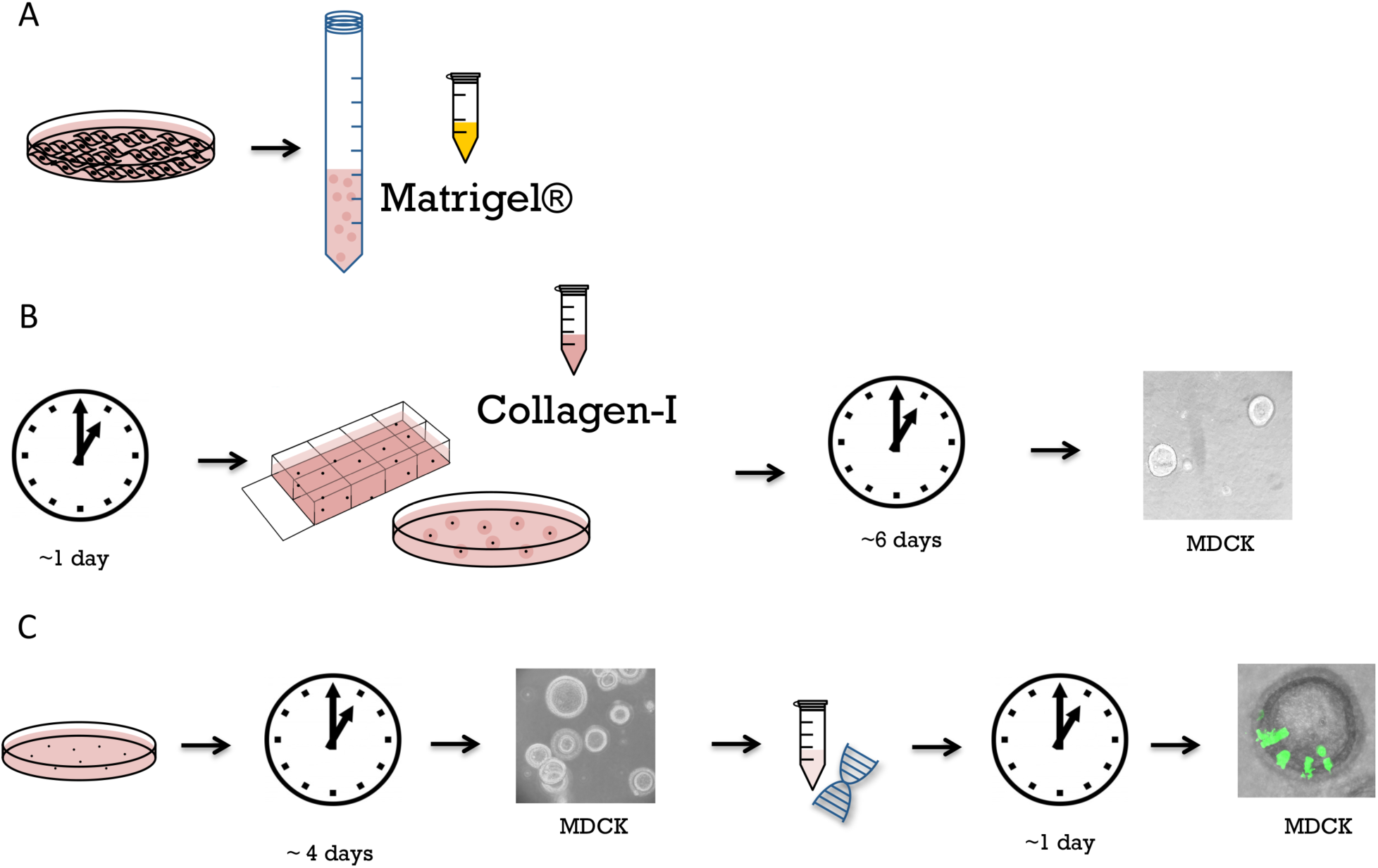
Preparation of polarized epithelial spheroid model. A. Overnight incubation of the cells with dilute Matrigel®. Cells are trypsinized, counted and incubated with the sub-gelation concentration of Matrigel® (1 mg / mL) overnight in a 15 mL conical tube. The incubation results in formation of cell clumps, which are later used as starter material for polarized epithelial spheroids. B. Incorporation of epithelial spheroids into hydrogels immediately after overnight incubation. Cell clumps are incorporated into collagen-I gels using either a sandwich system in a 8-chamber slide or 100 µL gel drops placed on the bottom of a 10 cm culture dish. Over the course of six days the spheroids grow and polarize. C. Growth of polarized epithelial spheroids in suspension. After overnight incubation with sub-gelation concentration of Matrigel® the cell clumps are transferred into a cell culture dish filled with culture media, in which they grow and polarize over the course of 4 days. The spheroids then can be transfected (optional) and seeded into hydrogel.

Cell aggregates can then be left to mature into spheroids in the Matrigel® suspension, or immediately incorporated into collagen-I hydrogels (Figure 1 B and C). For growth in suspension, the 1.5 mL of spheroid preparation can be transferred to a 35 mm dish containing 1 mL of warm culture media and incubated for 2 days at 37 °C under 5 % CO_2_. The suspension can then be transferred to a 60 mm dish containing 3 mL of warm culture media and incubated for additional 2-3 days.

### Incorporation of polarized epithelial spheroids into hydrogels

Collagen-I should be re-suspended in 0.02 N acetic acid at 3 mg / mL. This collagen stock solution can be aliquoted and stored at 4 C°. Combine the collagen solution with neutralizing solution (0.52 M Sodium Bicarbonate, 0.4 M HEPES and 0.08 N Sodium Hydroxide) and OPTI-MEM media at a ratio of 615 : 312 : 77 (collagen-I : OPTI-MEM: neutralizing solution) and keep the mixture on ice. To prepare to embed the spheroids, first coat the bottom of the wells of an 8-chamber slide with 50 µL of collagen solution to provide a layer of collagen-I on the surface; incubate the slides for 45 min at 37 °C to allow gelation.

Spheroids can be embedded into gels either after being grown to full polarity in the dilute Matrigel® solution or after the initial overnight incubation with Matrigel®. To embed the spheroids, transfer 1 mL of spheroid solution to a 15 mL conical tube, briefly spin it down and wash by gentle pipetting with 4 mL of cell culture media, briefly spin down again, and re-suspend in 400 µL of fresh cell culture media. As above, combine collagen-I stock solution (3 mg / mL in 0.02 N acetic acid) with neutralizing solution and spheroids in culture media in the ratio of 615 : 312 : 77. Pipette 50 µL of the collagen mixture containing cells onto the previously coated 8-chamber slide wells and gently spread it out into an even layer with the pipette tip; incubate for 45 min at 37 °C to allow gelation. Add 350 µL of fresh warm cell culture media to each well after the incubation and maintain the slides at 37 °C incubator with 5 % CO_2_. Change culture media every third day or as needed.

### Preparation of tumor spheroids

An overview of the epithelial spheroid model production protocol is depicted in Figure 2. Spheroids are prepared as described previously (McLane and Ligon, 2016). Briefly, MDA-MB-231 cells are harvested, trypsinized and counted. Aliquot 50,000 cells in 50 µL of culture media into wells of ultra-low attachment round bottom 96-well plates. Incubate plates at 37 °C under 5 % CO_2_ for 48 hours, which allows the cells coalesce into clumps. To further facilitate tumor spheroid formation, add 25 µL of growth factor-reduced Matrigel® (3 mg / mL in OPTI-MEM I) for a final concentration of 1 mg/mL, which is below the critical gelation concentration of Matrigel®. Incubate the coalesced masses for an additional 24 – 48 hours. After secondary incubation, cells will form tight tumor spheroids with a basement membrane mimetic external layer.

**Figure 2:**
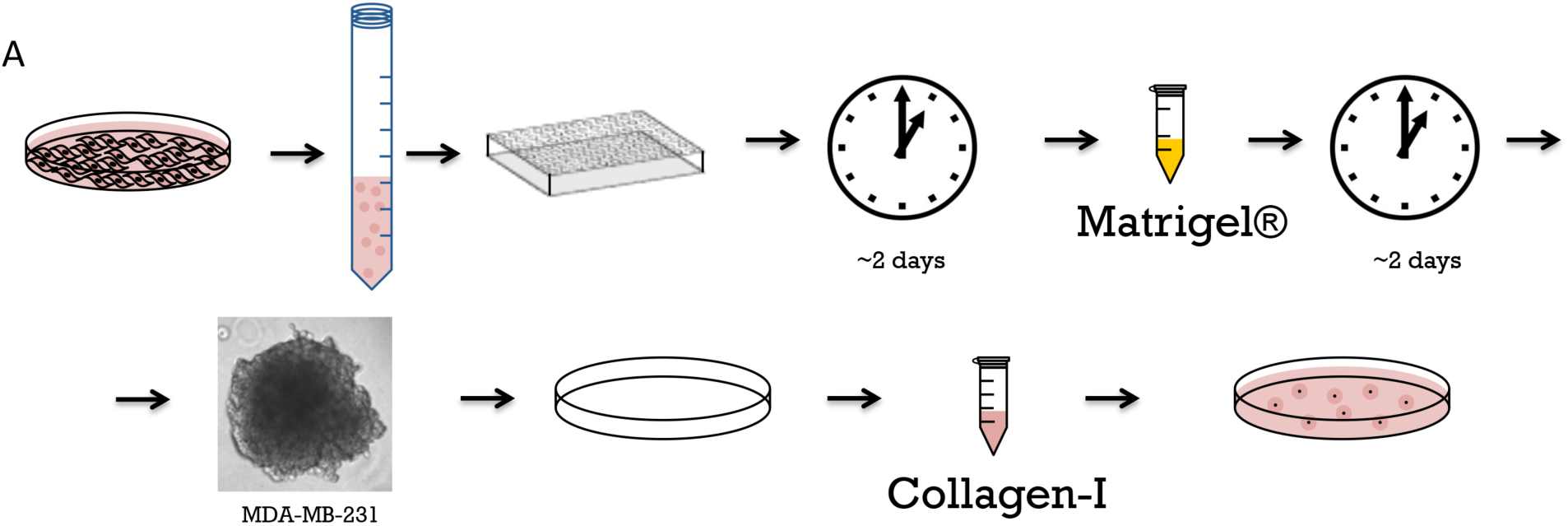
Preparation of tumor spheroid model. A. Tumor model preparation and seeding to collagen gel. Cells are trypsinized, counted and distributed into low-attachments 96-well plate, then incubated for two days, at which point cells coalesce. Sub-gelation concentration of Matrigel® (1 mg / mL) is then added to coalesced cells to provide basement membrane components and facilitate spheroid formation on the course of additional 48 hours. Formed spheroids are then incorporated into collagen gels and hydrogel droplets are placed in 10 cm cell culture dishes for further growth.

### Incorporation of tumor spheroids into hydrogels

Hydrogels are prepared as above. Briefly, collagen-I stock solution is combined with neutralizing solution and cell suspension media at a ratio of 615 : 312 : 77. Alternately, poly (ethylene glycol)-di (succinic acid *N*-hydroxysuccinimide ester) (PEG-diNHS) dissolved in DMSO (100 mg / mL, product E3257, Sigma-Aldrich, molecular weight 456.36) can be added to the gel to increase the gel stiffness by crosslinking the collagen fibers. In our hands, a ratio of 615 : 308 : 77 : 4 for collagen-I : suspension media: neutralizing solution: PEG-diNHS / DMSO resulted in a fourfold increase in gel stiffness (from ∼200 Pa to ∼800 Pa) (McLane and Ligon, 2015). These gels can also be pre-populated by stromal cells, such as fibroblasts by adding fibroblasts into the collagen solution prior to gelation (McLane and Ligon, 2016).

To make spheroid-containing hydrogels, transfer pre-formed spheroids in 2 μL media droplets to 10 cm dishes (eight spheroids per dish). Add 100 μL of collagen-I solution to each spheroid droplet, briefly mixing in the pipette tip and re-depositing in the dish. Incubate dishes for 45 min at 37 °C to allow gels to form, then add 10 mL culture media to the dish and release the hydrogels from the surface with a spatula. Culture all hydrogels on an orbital shaker to ensure they do not reattach to the culture vessel.

### Transfection of spheroids with plasmid DNA

To transfect cells in spheroids, allow them to form to the desired stage in suspension (e.g. grow epithelial spheroids to full polarity for 5 days). Wash 1 mL of spheroids in suspension as described above. Spin spheroids down gently to pellet and aspirate the media, then add 100 µL of SE Cell Line 4D-Nucleofector® X Kit and transfect with 4 µg of plasmid of choice. Here we used GFP plasmid (Clonetech, currently Takara Bio USA, Fremont, CA) diluted in MilliQ sterile-filtered water (Figure 1C). Transfect the cells using the 4D-Nucleofector protocol CA-152. After transfection, transfer the spheroids to a 35-mm dish with 2 mL of fresh warm media and leave spheroids for four hours to recover post-transfection. To form spheroid-containing hydrogels, collect the spheroids, briefly spin down and re-suspend in 400 µL of culture media. The spheroids then can be seeded in collagen gels as described above. A simplified illustration of 3D transfection workflow is shown in Figure 1C.

### Immunocytochemistry

The cells in the hydrogels can be stained for protein markers using conventional primary and secondary antibodies. The spheroids in gels are fixed with 4% paraformaldehyde in PBS for 45 min at 37°C, then permeabilized with 0.25% Triton-X in dH2O for 45 min at room temperature, washed briefly with PBS + 0.05% sodium azide (PBS-NaN_3_), and blocked for 2 hr at room temperature or overnight at 4°C in blocking solution (5% goat serum, 1% BSA, 0.05% NaN_3_ in PBS). Spheroids can then be incubated with primary antibodies diluted in PBS + 0.05% sodium azide overnight at 4°C. For that purpose, gels and diluted antibodies are placed in 1.5 mL conical tubes on a nutating shaker. Primary antibody incubation is followed by three PBS-NaN_3_ washes, 60 minutes each. Then, secondary antibodies and stains, such as DAPI or phalloidin diluted in PBS + 0.05% sodium azide are applied overnight at 4°C. The three 60 min washes with PBS-NaN_3_ are repeated after the overnight incubation. Fixed hydrogels with spheroids can be stored in 1.5 mL conical tubes in PBS-NaN_3_.

In the example shown here (Figure 3) the primary, antibodies used were: alpha tubulin @ 1: 500 (product T9026, Sigma-Aldrich, St. Louis, MO; Collagen-IV, product GTX26311, GeneTex, Irvine, CA). Secondary antibodies were used @ 1:300 (Alexa Fluor, Jackson labs, Bar Harbor, ME) together with rhodamine phalloidin for f-actin (product P1951, Sigma-Aldrich, St. Louis, MO) and DAPI (4’,6-diamidino-2-phenylindole).

**Figure 3:**
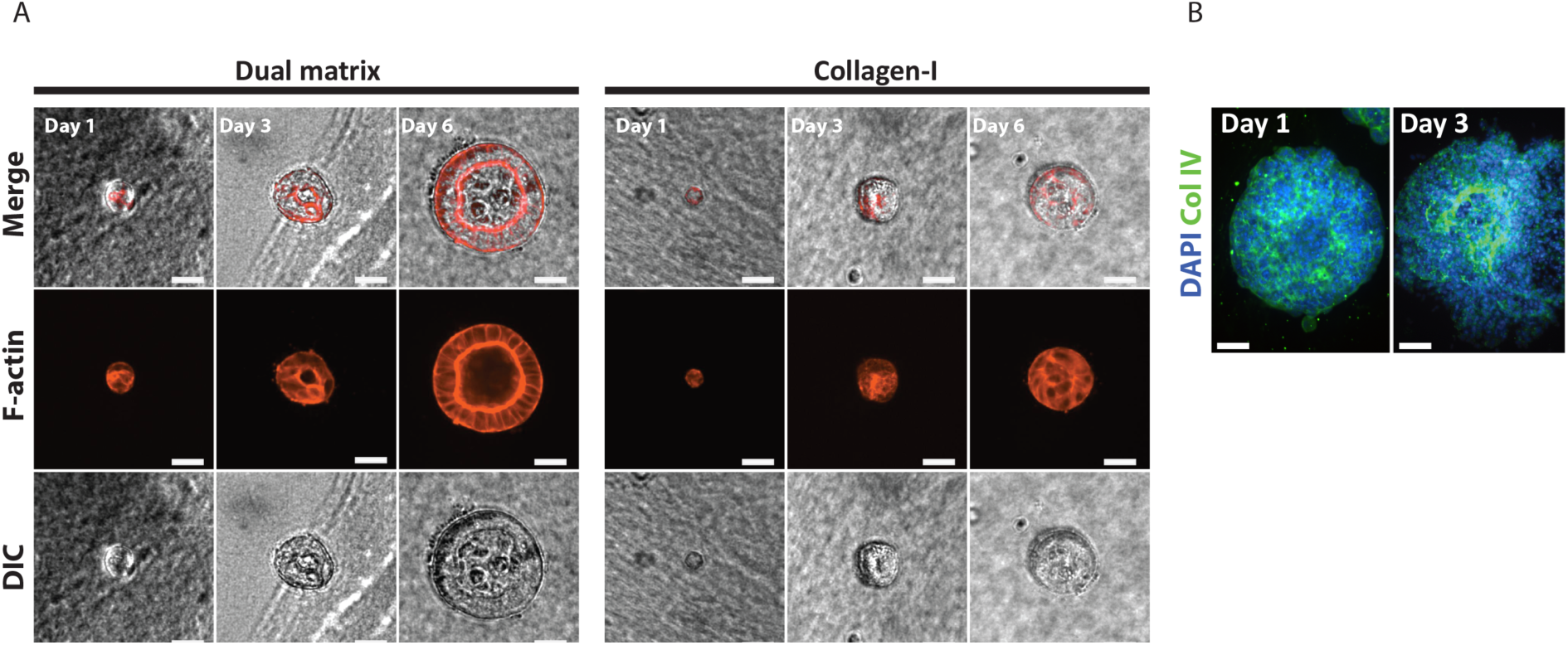
Polarized and tumor spheroids formation timeline. A. Comparison of speed of polarization of epithelial cells in dual-matrix versus collagen-I only culture. Spheroids are grown either using dual matrix system (left panel), or only in collagen-I (right panel). Spheroids are fixed on day 1, day 3 and day 6 in both systems and morphologies of spheroids are compared through immunostaining for filamentous actin (phalloidin, red). Examples of spheroids in both systems are shown for each day. Scale = 30 µm. B. Tumor spheroids of MDA-MB231 breast cancer cells grown in dual matrix system.

### Microscopy

The spheroids in the gels can be imaged at high magnification using both DIC and fluorescent microscopy. To prepare fixed cells for a imaging, place a gel in a 35 mm glass-bottom dish (MatTek), remove excess PBS with a Kimwipe and apply a drop of SlowFade® Diamond antifade mountant, allowing the gel to incorporate the antifade reagent for approximately 1 minute, and then placing a glass coverslip on top to flatten the gel, optionally adding a 1 g precision weight on top to further flatten the gel.

In the example shown here (Figure 3), imaging was done on an inverted microscope (DMI 4000B Inverted Microscope, LEICA Microsystems) outfitted with an ORCA-ER digital camera (Hamamatsu Photonics) and a Yokogawa spinning disc confocal using Volocity imaging software (Improvision/PerkinElmer).

### Cell isolation for nucleic acid or protein extraction

Protein and nucleic acid can be extracted from the cells grown in hydrogels for use in western blotting, PCR, and other application. To isolate cells, treat the gels with collagenase (product 02195109, MP Biomedicals) at 10 mg/mL until gels are digested (30 to 60 min). During digestion, place the tubes on a rotational shaker at 37 °C and monitor the tubes in five minute increments until the gels are completely digested. Centrifuge the digested gels at 300 xg for 5 minutes to pellet cells, and aspirate the digested collagen with a pipette. Wash the cell pellets twice with PBS by re-suspending cells in 1 mL PBS and repelleting cells with 300 xg spin. The cells can be then frozen, or used for nucleic acid or protein extraction using standard protocols.

## RESULTS AND DISCUSSION

Here, we present a method to generate a model epithelial tissue, in which the organization, composition and physical properties of the ECM are physiologically appropriate, composition is controlled, and stiffness can be tuned (McLane and Ligon, 2016). This model system can be used to recapitulate normal epithelial organization, or an early *in situ* tumor. In both cases, the cells are encased in a basement membrane, initially nucleated by Matrigel®, and then are surrounded by a stiffness-controlled stromal matrix, composed of type I collagen, in which stromal cells such as fibroblasts can also be embedded. Other ECM components can be added in to the stromal mixture as well to increase the physiological accuracy.

### Formation of normal polarized epithelial spheroids

The spheroids formed with the dual matrix method show early and robust polarization. As shown in Fig. 3, spheroids at day 1 are composed of multiple cells, distinctly visible with F-actin labeling (red). By day 3, the cells in spheroids have begun to assume the columnar morphology characteristic of polarized epithelial cells, and the spheroid has also begun to establish a hollow core. By day 6, the cells in the spheroids show distinct polarized morphology, and the hollow core is fully formed. In comparison, single cells seeded in a collagen-I gel form a mostly disorganized clump of cells by day 3, and do not show signs of polarization (columnar cell morphology, hollow core formation) by day 6. This side-by-side comparison clearly illustrates the increased speed of polarization and spheroid formation in the dual-matrix system as compared with a single matrix collagen-I system.

### Formation of tumor spheroids

Tumor cell spheroids formed with the dual matrix method demonstrate behaviors characteristic of a tumor *in-situ*, such as matrix invasion, while spheroids formed of cells of non-metastatic lineage do not (McLane and Ligon, 2016). This normalized behavior from non-metastatic cells, the expected original hypothesis, is not what has been historically observed in single matrix culture models and is apparently mediated by the establishment of a basement membrane prior to hydrogel incorporation. This clearly illustrates the importance of the dual matrix system and the ability to mimic the *in-vivo* microenvironment in comparison to single matrix systems.

### Comparison with other methods

For over 30 years, Collagen-I hydrogels have been used to model the three-dimensional cellular environment. Numerous other 3D culture methods have been developed as well (Kimlin et al., 2013), including, but not limited to, cell culture upon or within natural protein hydrogels of Matrigel®, fibrin, hyaluronic acid (Masters et al., 2004), chitosan (Azab et al., 2006), and alginate (Barralet et al., 2005) as well as non-biological substrates or hydrogels including polyvinyl alcohol (Martens, 2000), poly-L-lactic acid (McLane et al., 2014a; Wang et al., 2009), and polyethylene glycol (Sawhney et al., 1993). Many of these hydrogel types have been combined and/or chemically modified to gain specific structural or mechanical characteristics, such as pore size, fibril size, alignment, stiffness (Chenite et al., 2001; Munoz-Pinto et al., 2012; Roeder et al., 2002; Wang et al., 2010; Deng et al., 2010; McKay et al., 2014; Wang and Stegemann, 2011; Liang et al., 2011), or to modulate ligand availability (Liu et al., 2010; Yoshida et al., 1997; Krause et al., 2008; Swamydas et al., 2010; Gandavarapu et al., 2014). The method described here utilizes several of these hydrogel strategies to recapitulate two distinct characteristics of the epithelial microenvironment – the dual matrix type and tissue organization of a normal epithelial tissue or a mammary tumor *in situ*, as well as the altered mechanical properties of the tumor-associated stroma.

The stiffness of a collagen-I hydrogel can be modulated by varying parameters such as pH, gelation temperature and collagen concentration (Li et al., 2009; Yang et al., 2009; Plant et al., 2009), as well as by the incorporation of other biotic or abiotic materials (Wang and Stegemann, 2010; Ulrich et al., 2010; Batorsky et al., 2005; Li et al., 2012; Ulrich et al., 2011; Song et al., 2010; Deng et al., 2010; Krause et al., 2008). However, altering these parameters or incorporating a secondary material alters the structure of the hydrogel and/or availability of the collagen-I ligand. To avoid these potential changes, we control collagen-I hydrogel stiffness independently of pH, temperature and protein concentration and without the incorporation of a second material by crosslinking the collagen-I with poly-(ethylene glycol)-di (succinic acid N-hydroxysuccinimide ester) (PEG-diNHS) (Abdella et al., 1979). PEG-diNHS makes short crosslinks between proteins by forming amide bonds between the collagen-I and itself to tether collagen molecules together, which mimics cross-links formed in vivo (Wallace, 2003). These collagen-I PEG-diNHS hydrogels have been previously used in studies of tumor spheroid formation and in tissue engineering, and show good biocompatibility (Jeong et al., 2013; McLane and Ligon, 2015a; Liang et al., 2011). These hydrogels also have defined, reproducible matrix stiffness within the range of epithelial and mammary physiology (Levental et al., 2009; Paszek et al., 2005), in contrast to many studies of cell-matrix interaction which either greatly exceed the physiological range (Leight et al., 2012; Tilghman et al., 2010; Pathak and Kumar, 2012; Chia et al., 2012), or do not measure or consider the stiffness of their model system.

The basement membrane (BM) extracellular matrix is made up of different proteins from that of connective tissue, and those that are shared between the two are present in different concentrations (Shoulders and Raines, 2009). In the case of normal epithelial tissue or a carcinoma *in situ*, an early stage in tumor development in which the BM is still intact, the cells are surrounded by the BM. Tumor cells must degrade this BM or otherwise circumvent it before invading into the stromal tissue. Our model of both normal epithelial spheroids and of a tumor *in situ* utilizes Matrigel®, a commercially available sarcoma produced protein mixture rich in basement membrane proteins (Hughes et al., 2010), to jumpstart BM formation. To accomplish this, we form cell aggregates or spheroids in the presence of dilute Matrigel®. We use Matrigel® at a concentration below the critical gelation concentration, so it does not form a gel, but allows basement membrane components to be adsorbed to the surface of the cells during aggregate or spheroid formation. Most epithelial cells will also secrete basement membrane proteins and form a basement membrane by themselves, but this process can take over a week. By providing building blocks, we can significantly accelerate basement membrane formation. We then incorporate the BM-coated spheroid into a stromal mimetic collagen-I hydrogel. This creates an *in vivo-*like organization and matrix composition in which epithelial cells are encased in a BM surrounded by a stromal matrix and stromal cells. The organization of the model allows for the transmission of both chemical and mechanical signals between epithelial cells and the stromal matrix and any stromal cells incorporated in it. There are few other experimental systems currently in use which embed basement membrane coated spheroids into collagen-I stromal matrices of defined stiffness, although there are co-culture systems which allow for chemical signaling between cell types (Peng et al., 2013; Bischel et al., 2015) or which investigate cell-cell or cell-matrix interactions with multiple matrices, although the organization of the matrices may not be as physiologically relevant as ours (Viney et al., 2009; Krause et al., 2008; Swamydas et al., 2010).

The most common 3D hydrogel based approach to investigate tumor cell invasiveness involves starting with a single cell suspension in a hydrogel and then allowing the cells to proliferate to form acinar structures (Chambers et al., 2011; Krause et al., 2008; Swamydas et al., 2010; Liang et al., 2011). Our model departs from this method by first forming epithelial tumor spheroids in the presence of basement membrane components to accelerate the formation of a basement membrane. These formed spheroids are then incorporated into the stromal matrix after a coherent structure in which the cells have developed significant apical-basal polarity has formed. We believe this is more representative of a tumor *in situ* and will yield more translational results as tumors develop from existing tissue, not from single cells within a matrix. We have recently used this method to show that spheroids of both normal MCF10A cells and more metastatic MDA-MB-231 cells behave somewhat differently than when grown in a less physiologically relevant system (McLane and Ligon, 2016). For example, it has previously been suggested that the phenotypically normal MCF10As become invasive when grown in a stiff matrix, but we showed that when the MCF10As are grown in this physiologically appropriate two matrix system, they do not show an invasive phenotype with increased stromal stiffness.

Similar to the methods described above to investigate the tumor microenvironment, studies of normal epithelial biology in 3D have also typically started from single cells seeded in a collagen-I matrix, and then allowed to develop for 10-12 days into mature cysts (Montesano 1991, reviewed in Zegers 2003, Belmonte 2008). In other studies, cells were seeded in Matrigel® instead (Belmonte 2008). In both cases, the organization of the model did not fully recapitulate normal tissue arrangement. In addition, another drawback to this approach is that during the long incubation necessary to achieve fully polarized spheroids, some cells can migrate away from the spheroid to the edge of the gel, where they form a 2D monolayer that can interfere with imaging, and perhaps alter the mechanical properties of the matrix.

We have found that growing cells in a sub-gelation concentration of Matrigel® prior to seeding them in collagen-I promotes the formation of small multi-cell clusters (nucleated aggregates). Seeded into the collagen gel, these starter spheroids develop into mature spheroids in ∼5-6 days, thus shortening the experiment preparation time by ∼60% or up to 6 days. Alternatively, spheroids can be grown to full polarization in the dilute Matrigel® solution before incorporation into the collagen-I gel, which further increases the maturation speed, with most spheroids ready to use by day 4.

Another major issue in 3D cell culture is that it is difficult to perform genetic manipulations on cells that are encapsulated in a hydrogel. One way around this limitation is to create stably transfected cell lines with a drug-inducible construct. Here we have developed methods to manipulate cells while they are growing in the dilute Matrigel® solution via methods such as electroporation (Nucleofection™), lipid-based reagents or iron-oxide nanoparticles (Magnetofection™). We have recently used this model system to investigate the mechanisms of epithelial morphogenesis and have shown that spheroids grown with this method display the same markers of polarity and respond to growth factor stimulation in the same way as the traditional spheroids grown in collagen-I (Bogorodskaya and Ligon, *submitted*).

### Controls and caveats

While we discuss the basics of spheroid formation and collagen-I hydrogels, it is important to note that there are a large number of parameters that can affect the properties of the model system. Altering a parameter can drastically change the collagen-I fiber size, hydrogel porosity, mechanical properties, spheroid size, spheroid number and cell survival.

As with any culture system, selection of cell culture media is critical and the effects of different media on all cell types used in the system must be evaluated. Cross-linkers can cause viability issues with some cell types, so we also recommend evaluating the viability of your cells after incorporation into the PEG-diNHS cross-linked collagen-I stromal hydrogel. Finally, there are many opportunities for variation in preparing the various reagents used in making the hydrogels, so we also recommend evaluating the stiffness of the collagen-I hydrogels to ensure that they are of desired stiffness. We have done so via bulk rheometry, but other methods such as extensiometry (Drury et al., 2004), nano-indentation by Atomic Force Microscopy (Soofi et al., 2009), or even embedded magnetic particles (Chippada et al., 2009) could be used as well.

### Limitations

The initial size of spheroids may be potentially limiting for some experimental scenarios. Although it is possible to generate very large spheroids, for this method, the spheroid must fit through the opening of micropipette tips (we use 200 μL tips for all of our spheroid handling) and scale to the volume of the tip. This limitation is however easily overcome by using larger pipettes and larger volumes of hydrogel. Although we have observed no nutrient limitation or waste product induced cell death at the scale we have used (spheroids up to ∼2 mm^3^), these are potential concerns for larger spheroids. For polarized cysts, the size of the resulting cyst will be dependent on the initial starter spheroid, and high variability of cyst sizes in possible, with spheroids ranging from 100 µm to 500 µm and larger in diameter. For cysts left to mature in Matrigel® suspension, most of the cysts will be ready on day 4, but those left in Matrigel® will continue increasing in size, reaching up to 500 µm in diameter.

## ACKNOWLEDGEMENTS

We would like to thank Mariah Hahn and Ryan Gilbert as well as members of their labs, and former Ligon lab member Joseph Wiegartner for their assistance with biomaterials. This work has been supported by the American Cancer Society Research Scholar Grant (RSG-10-245-01-CSM) and by the National Institutes of Health grant R01GM098619.

